# Privacy-preserving biomedical database queries with optimal privacy-utility trade-offs

**DOI:** 10.1101/2020.01.16.909010

**Authors:** Hyunghoon Cho, Sean Simmons, Ryan Kim, Bonnie Berger

**Affiliations:** Broad Institute of MIT and Harvard; Harvard University; Computer Science and AI Laboratory, MIT; Department of Mathematics, MIT

## Abstract

Sharing data across research groups is an essential driver of biomedical research. In particular, biomedical databases with interactive query-answering systems allow users to retrieve information from the database using restricted types of queries (e.g. number of subjects satisfying certain criteria). While these systems aim to facilitate the sharing of aggregate biomedical insights without divulging sensitive individual-level data, they can still leak private information about the individuals in the database through the query answers. Existing strategies to mitigate such risks either provide insufficient levels of privacy or greatly diminish the usefulness of the database. Here, we draw upon recent advances in differential privacy to introduce privacy-preserving query-answering mechanisms for biomedical databases that provably maximize the expected utility of the system while achieving formal privacy guarantees. We demonstrate the accuracy improvement of our methods over existing approaches for a range of use cases, including count, membership, and association queries. Notably, our new theoretical results extend the proof of optimality of the underlying mechanism, previously known only for count queries with symmetric utility functions, to asymmetric utility functions needed for count queries in cohort discovery workflows as well as membership queries—a core functionality of the Beacon Project recently launched by the Global Alliance for Genomics and Health (GA4GH). Our work presents a path towards biomedical query-answering systems that achieve the best privacy-utility trade-offs permitted by the theory of differential privacy.

## 1 Introduction

The fast accumulation of biomedical datasets around the globe, including personal genomes and medical records, hold immense potential to advance biology and medicine. However, most of these data are held in isolated data repositories (e.g. different hospitals or biobanks); sharing this data across repositories is often infeasible due to data privacy concerns. A pressing challenge in the biomedical community is to develop systems that allow researchers to jointly leverage large data collections across multiple sites in order to gain more accurate and refined biomedical insights crucial to realizing the vision of personalized health.

In response to this need, several large-scale databases in both medical and genomics communities have developed interactive query-answering systems in order to allow external researchers and clinicians to utilize their database in a limited and controlled fashion [1–6]. For example, medical data repositories, such as i2b2 and STRIDE [4, 5], allow researchers designing clinical studies to query how many patients in the database satisfy a given set of criteria prior to requesting for data access or recruiting patients, a workflow commonly known as *cohort discovery*. In addition, recently emerging genomic “beacon” services [6] allow users to query whether or not a given genetic variant is observed in the database, a workflow we refer to as *variant lookup*. The Beacon Project by the Global Alliance for Genomics and Health (GA4GH) [7] has helped launch a network of over 100 beacons to date around the world [6]. These interactive systems are poised to play a key role in driving data sharing efforts in the biomedical community.

Yet despite the limited scope of allowed queries and the fact that only aggregate-level information is shared, query-answering systems can still leak sensitive information about the underlying individuals [8–11]. One could, for example, ask for the number of 25-year-old females on a certain medication who do not have a particular disease. If the answer returned is zero, we know that any 25-year-old female patient in the database who is on that medication has that disease, a fact the patients might wish to keep private. Moreover, given access to an individual’s genotype, a small number of queries to a beacon server would be sufficient to reveal whether the individual is included in the database [10, 11]. This information could potentially be detrimental to the individual if the underlying cohort represents a group of individuals with sensitive characteristics (e.g. a rare medical condition).

To address these privacy concerns, existing systems and studies have attempted to improve individual privacy in query-answering systems by perturbing query results with a small amount of noise in order to reduce sensitivity to underlying individuals [11, 12]. However, existing efforts either lack rigorous theoretical guarantees of privacy or introduce an excessive amount of noise into the system, limiting their effectiveness in practice. For instance, i2b2 and STRIDE add truncated Gaussian noise to the number of subjects matching a given filter without consideration of formal models of privacy. Recent studies [9,13] have proposed to apply the theoretical framework of differential privacy [14, 15] to the cohort discovery problem. *As we demonstrate in our work, existing methods perturb the results more than is necessary to achieve a desired level of privacy.* For beacon servers, although real-world systems are yet to adopt a standard protocol to remedy privacy risks, Raisaro et al. [11] recently explored potential attacks on beacon servers and provided a set of risk mitigation strategies. Unfortunately, their techniques are based upon a simplified model of genotype distribution, which is prone to sophisticated attacks that exploit the data patterns not captured by the model (e.g. deviations from Hardy-Weinberg equilibrium), as we show in our results.

Here, we build upon recent advances in differential privacy (DP) to introduce query-answering systems with formal privacy guarantees while ensuring that the query results are as accurate as theoretically possible. We newly propose to leverage the truncated *α*-geometric mechanism (*α*-TGM), previously developed for a limited class of count queries in a theoretical context [16], to limit disclosure risk in both cohort discovery and variant lookup problems in biomedical databases. We show that *α*-TGM, combined with a post-processing step performed by the user, provably maximizes the expected utility (encompassing accuracy) of the system for a broad range of user-defined notions of utility and for any desired level of privacy. Notably, the optimality of *α*-TGM was previously known for only symmetric utility functions, which are insufficient for workflows we typically encounter in biomedical databases [9]. We newly extend this result to a more general class of utility functions including *asymmetric* functions, *thereby answering an open question posed in the original publication of α-TGM [16]*. Asymmetric utility is a desirable notion in cohort discovery applications where overestimation is often more desirable than underestimation [9]. Moreover, we demonstrate that *α*-TGM can be transformed to obtain an optimal DP mechanism for the variant lookup problem, a novel result enabled by our optimality proof for generalized utility.

In addition to our theoretical results, we empirically demonstrate the accuracy improvements of our proposed optimal DP mechanisms for cohort discovery, variant lookup, as well as for chi-squared association tests built upon count queries. We also newly illustrate how a user’s ability to choose a prior belief over the true answer can further boost the accuracy of the query results in our DP framework, without incurring additional privacy cost. Given the rapidly-growing availability of interactive biomedical databases around the world, principled approaches to mitigating their privacy risks are becoming increasingly essential. Our work shows how one can leverage the theory of differential privacy to protect the privacy of individuals in these systems, while simultaneously maximizing the benefits of data sharing for science.

A MATLAB implementation of our DP mechanisms and scripts for reproducing the results presented in this work are available at: https://github.com/hhcho/priv-query.

## 2 Definitions and Review of Existing Methods

### 2.1 Review of Differential Privacy

Differential privacy (DP) [14, 15] is a theoretical framework for sharing aggregate-level information about a dataset while limiting the leakage of private information about individuals in the dataset. The idea behind differential privacy is that, if we have a dataset from which we want to release some statistic, it should be difficult to tell the difference between the statistic calculated on our dataset and the statistic calculated on a dataset that differs from ours by exactly one individual. Differential privacy achieves this property by adding a controlled amount of statistical noise to the shared data in such a way that it ensures the statistical impact of any particular individual in the database is smaller than the desired level. Originally developed in cryptography, the theory of differential privacy has found many applications and theoretical connections in diverse domains including privacy-preserving machine learning [17], robust statistics [18], and quantum computing [19], and has recently been adopted to protect private individuals in real-world systems, e.g. by Google and the United States Census Bureau [20].

More formally, assume that we have a dataset, denoted *X*, and we want to release some statistic, denoted *f* (*X*), that has been calculated on our data set. This statistic, however, may not preserve the privacy of the individuals in *X*. As such, we will instead release a perturbed statistic, denoted *F* (*X*), which approximates *f* (*X*) while still achieving a certain level of privacy. This level of privacy is measured by a parameter *ϵ* > 0, where the closer to zero *ϵ* is, the more privacy is retained. The goal is that, for every pair (*X, X*′) of “neighboring” datasets (i.e., *X* and *X*′ are of the same size and differ by exactly one individual), we have that for any possible outcome *y* in the image of *F*,

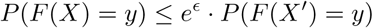

Any *F* that satisfies this property is said to be *ϵ-differentially private*. Intuitively, this property ensures that it is statistically hard to distinguish *F* (*X*) from *F* (*X*′), thereby ensuring that no one individual loses too much privacy when *F* (*X*) is released.

One can extend this framework to protect multiple queries using the following composition properties: Given a sequence of *k* statistics *F* = (*F*_1_, …, *F*_*k*_) where each statistic *F*_*i*_ is *ϵ*_*i*_-differentially private, the overall algorithm *F* is 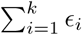-differentially private (*sequential composition*). When each of the statistics in *F* is computed based on a disjoint subset of the individuals in the dataset, *F* is max_*i*∈{1,…,*k*}_ *ϵ*_*i*_-differentially private (*parallel composition*). More advanced composition techniques with tighter privacy bounds on the overall mechanism have been proposed [21]. These tools enable database owners to assign a privacy “budget” to each individual user and keep track of the combined privacy level *ϵ* throughout the user’s interaction with the database, which reflects how much private information is revealed to the user overall.

### 2.2 Previous Methods for Cohort Discovery and Variant Lookup with Privacy

Key use cases of biomedical query-answering systems include *cohort discovery* and *variant lookup*. In cohort discovery, clinical researchers ask how many patients in the medical record database meet a certain condition. This information can help researchers to design studies and assess their feasibility without having to obtain sensitive patient data first. In variant lookup, researchers or physicians ask whether a given genetic variant is observed in the individuals in the database. This query can facilitate the matching of patients with similar genetic determinants of disease across different hospitals, or improve the phenotypic characterization of genetic variants of interest. Here we review existing proposals for mitigating privacy risks associated with these two types of queries.

#### Cohort discovery with privacy

Existing cohort discovery systems such as i2b2 and STRIDE [4, 5] use approaches to protecting privacy that give no real privacy guarantees, whereby they release perturbed counts for some users instead of raw counts. Both of these methods work by adding truncated Gaussian noise to the query results, which unfortunately do not provide formal privacy guarantees. In order to remedy this issue, Vinterbo et al. [9] suggested a method to produce differentially private answers to count queries. Although their work was not the first attempt to use differential privacy in a medical research context, it is the first we are aware of to do so as a way to improve medical count queries [22–25].

We briefly review the approach of Vinterbo et al. [9] here. In a nutshell, the authors assume that there is a database consisting of *n* patients, and they want to know the number of people in that database meeting a certain condition. To accomplish this goal, they introduce a loss function *ℓ* defined by

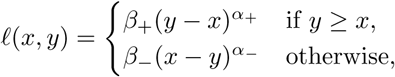

where *α*_+_, *α*_−_, *β*_+_, *β*_−_ are parameters given by the user, *y* is an integer in the range [*r*_min_, *r*_max_] to be released by the mechanism, and *x* is the true count. This loss function measures the loss of approximating *x* with *y*.

Note that *ℓ* has sensitivity (i.e., maximum change caused by substituting a single individual) Δ*ℓ* = max(*β*_+_, *β*_−_). Therefore, if we define a random function *X*_*ω*_ such that, for a given value *y, P* (*X*_*ω*_(*y*) = *x*) is proportional to exp{−*ωℓ*(*x, y*)} for all integers *c* ∈ [*r*_min_, *r*_max_], then *X*_*ω*_ is 2Δ*ℓω*-differentially private. This approach is commonly known as the *exponential mechanism* [26]. *Given the above parameters and ϵ*, Vinterbo et al.’s mechanism [9] returns an estimate of *y* given by *X*_*ω*_(*y*), where *ω* = *ϵ*/(2Δ*ℓ*). Note that this result is *ϵ*-differentially private by the standard analysis of the exponential mechanism [26], though a more thorough analysis can show that this is not a tight bound on *ϵ* (see the Appendix).

A recently-proposed framework called MedCo [13] allows multiple hospitals to collectively answer cohort discovery queries without sharing the underlying databases. MedCo uses the *Laplace mechanism* to achieve *ϵ*-differential privacy, whereby the system adds noise from the Laplacian distribution with scale parameter 1/*ϵ* to the true count, rounds it to the nearest integer, then releases the result to the user. Unlike Vinterbo et al.’s approach, this framework does not allow users to choose their own notion of utility; instead, it approximately corresponds to a loss with *β*_+_ = *β*_−_ and *α*_+_ = *α*_−_ = 1.

#### Variant lookup with privacy

Shringarpure and Bustamante [10] were the first to demonstrate a *membership inference attack* on genomic beacon services, which we refer to as the SB attack. In their attack scenario, it is assumed that the attacker has access to (a subset of) the target individual’s genotypes. The attacker repeatedly queries the beacon with the individual’s data to infer whether or not the individual is included in the database. More precisely, the likelihood ratio over the query responses between the case where the database includes the individual and the case where it does not is used to statistically test whether the individual is indeed in the database. Depending on how rare the queried variants are, it has been shown that a small number of queries are sufficient for the SB attack to succeed.

Recently, Raisaro et al. [11] further explored this privacy threat for emerging beacon services, and proposed a set of risk mitigation strategies, which include: (i) only answering yes when the query variant is observed at least twice; (ii) answering incorrectly with probability *ϵ* for rare variants; and (iii) keeping track of the cumulative likelihood ratio in the SB attack for each individual and suppressing a user once a threshold has been reached. However, these strategies were specifically designed for and analyzed based on the SB attack, which assumes certain properties about the data distribution, e.g. that every variant satisfies Hardy-Weinberg equilibrium. As a result, although Raisaro et al.’s strategies are reasonable first steps, they do not guard against more sophisticated attacks that exploit data patterns not captured by the underlying model, as we demonstrate in our results. Differential privacy techniques, on the other hand, are agnostic to how the individual-level data is distributed and thus enable more rigorous guarantees of privacy.

## 3 Our Approach: Utility-Maximizing Differential Privacy Mechanisms

We introduce differential privacy (DP) mechanisms for cohort discovery and variant lookup problems that achieve provably optimal trade-offs between privacy and utility. Our approaches build upon the truncated *α*-geometric mechanism for the count query problem [16], which is well-studied in a theoretical context.

### 3.1 Count Queries for Cohort Discovery

Here we briefly describe previous work [16] and then turn to our generalizations of it.

#### *α*-geometric mechanism

Let *x* ∈ {0} ∪ ℤ+ be the true result of a count query. The *α*-geometric mechanism (for *α* ∈ (0, 1)) takes *x* as input and releases a perturbed value *y* = *x* + Δ ∈ ℤ, where

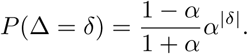

This mechanism achieves ln (1/*α*)-differential privacy, because changing a single database entry changes *x* by at most one; the likelihood of any observed value of *y* differs by at most a multiplicative factor of 1/*α* = *e*^ln(1/*α*)^ for neighboring *x* and *x*′ that differ by one.

#### Truncated *α*-geometric mechanism (*α*-TGM)

Now consider a mechanism that takes the output *y* from the *α*-geometric mechanism and truncates it to the range of possible results for a count query given a database of *n* entries to release *z* = min{max{0, *y*}, *n*}. This is called the *truncated α*-geometric mechanism. Due to the post-processing property of differential privacy, which states that any additional computation on differentially private statistics does not cause additional privacy loss, this mechanism is also ln(1/*α*)-differentially private.

#### Optimality of *α*-TGM for count queries with symmetric utility

Let *ℓ*(*x, y*) be a loss function that captures the disutility (i.e., how undesirable an outcome is to the user) of releasing a result *y* when the true result of the count query is *x*. We are interested finding a *ϵ*-DP mechanism that maximizes (minimizes) the expected utility (disutility) for any given *ϵ*. Note that we can parameterize the DP mechanism by the conditional probability distribution *q*(*y*|*x*). Formally, the utility-maximization problem is given by

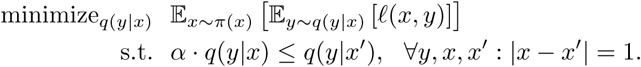

where *α* = *e*^−*ϵ*^ and *π*(*x*) denotes the prior belief over the true count *x*{0, …, *n*}.

The following theorem summarizes the main result of Ghosh et al. [16]:

##### Theorem 1

(Ghosh et al. [16]; adapted). *Given any prior belief π*(*x*) *and a symmetric loss function ℓ*(*x, y*) = *f* (|*x*−*y*|) *for a monotonically increasing f, a utility-maximizing ϵ-DP mechanism for the count query problem is obtained by applying a* (*π, ℓ*)*-dependent post-processing to the output of* exp(−*ϵ*)*-TGM.*

This is a striking result in that *the core component of the optimal mechanism (i.e., α-TGM) is agnostic to the user’s chosen prior distribution and loss function.* This theorem suggests that given a desired privacy level *ϵ*, the user can obtain a generic query response from the database based on the *α*-TGM and locally apply post-processing to obtain a tailored response that is provably optimal in terms of utility among all *ϵ*-DP mechanisms for this task. Furthermore, this scheme allows the user to locally try out different choices of *π* and *ℓ* without consuming additional privacy budget (which roughly corresponds to the number of queries a user is permitted). Thus, the theoretical results of Ghosh et al. [16] immediately provide an improved DP mechanism for cohort discovery compared to existing proposals [9, 13].

The optimal post-processing step is given by minimizing the expected loss under the *posterior* belief over the true count, conditioned on the output of the *α*-TGM. Formally, let *q*_*TGM*_ (*z*|*x*; *α, n*) be the conditional output distribution of *α*-TGM for a true count *x* based on a dataset of size *n*. Conditioned on the value of *z*, the posterior belief over *x* can be expressed as:

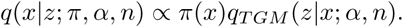

Next, define a map *T* : [*n*] ↦ [*n*] as

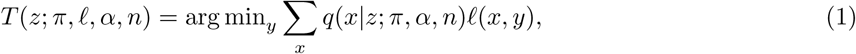

which represents the loss-minimizing guess for *x* conditioned on *z*. Finally, the *ϵ*-DP query response with maximum expected utility with respect to (*π, ℓ*) is given by *T* (*z*; *π, ℓ*, exp(−*ϵ*), *n*), where *z* is the output of the exp(−*ϵ*)-TGM.

#### Our contribution: extending the optimality of *α*-TGM to more general utility functions

A key limitation of Ghosh et al.’s scheme [16] is that Theorem 1 applies to only loss functions that are symmetric around the true query result. It has been previously noted that *asymmetric* utility functions are useful in cohort discovery [9]. For example, overestimating the amount of resources required to perform a clinical study due to overestimating the number of individuals who can be enrolled is more favorable than underestimating it only to encounter a resource bottleneck during the study.

To this end, we newly generalize Theorem 1 to a more general class of utility functions, notably including both symmetric and asymmetric functions, thereby broadly enabling the use of *α*-TGM for cohort discovery applications. In fact, Ghosh et al. posed the case of asymmetric utility as an open question in their publication [16], which we resolve in our work. Note that we generalize the utility even further to allow a different choice of utility for every possible value of the true count *x*, a result that we will return to in the following section on variant lookup. Our generalized theorem is as follows.

##### Theorem 2.

*Given any prior belief π*(*x*) *and a loss function ℓ*(*x, y*) = *f*_*x*_(*y* − *x*), *where f*_*x*_ *is quasiconvex and attains its minimum at zero for all x, a utility-maximizing ϵ-DP mechanism for the count query problem is obtained by applying a* (*π, ℓ*)*-dependent post-processing to the output of* exp(−*ϵ*)*-TGM.*

The proof is included in the Appendix. Note that the (*π, ℓ*)-dependent post-processing step is identical to the scheme we previously described for the symmetric case, except with expanded support for a broader range of loss functions the user can choose from.

### 3.2 Membership Queries for Variant Lookup

We newly show that *α*-TGM can be used to obtain an optimal DP mechanism for the variant lookup problem.

#### Our optimal differential privacy mechanism for variant lookup based on *α*-TGM

Formally, the problem can be described as follows. Let *X* be a dataset of *n* individuals represented as *X* = (*x*_1_, …, *x*_*n*_) ∈ 𝒳^*n*^. Given a user-provided predicate *q* : 𝒳 ↦ {0, 1}, we define *membership query* as the task of calculating

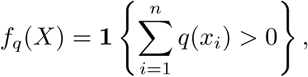

where **1**{·}denotes the indicator function. Our goal is to answer queries of this form while preserving the privacy of individuals. Here we will restrict our attention to stochastic mechanisms that consider the total count 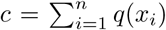 as input and probabilistically output a binary answer for the query. However, analogous to the results of Ghosh et al. [16] for count query, it can be shown this restriction is without loss of generality; i.e., our mechanism is optimal even among mechanisms that may depend on more fine-grain data patterns other than the count.

Akin to the count query setting, in order to quantify the usefulness of different DP mechanisms for membership queries, we introduce a loss function *ℓ*(*c, b*), representing how undesirable it is to output an answer *b* ∈ {0, 1} given a true count *c* ∈ {0, …, *n*}, as well as a prior belief *π*(*c*) over the true count.

#### Our main result

The following theorem summarizes our main result that the optimal DP mechanism for membership queries, which achieves the minimum expected loss with respect to a particular choice of *π* and *ℓ*, is obtained by a generic application of *α*-TGM followed by a local post-processing step by the user; only the latter step depends on *π* and *ℓ*.

##### Theorem 3.

*Given any prior belief π*(*c*) *and a loss function ℓ*(*c, b*) *satisfying ℓ*(0, 0) ≤ *ℓ*(0, 1) *and ℓ*(*c*, 0) ≥ *ℓ*(*c*, 1) *for c* > 0, *a utility-maximizing ϵ-DP mechanism for the membership query problem is obtained by applying a* (*π, ℓ*)*-dependent post-processing step to the output of* exp(−*ϵ*)*-TGM.*

The proof is provided in the Appendix. The optimal post-processing step of the above theorem proceeds as follows. Given a specific choice of *π* and *ℓ*, the user first transforms *ℓ* into the corresponding loss function *ℓ*′ in the count query setting as

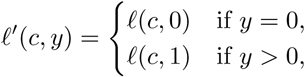

for *c, y* ∈ {0, …, *n*}. Let *z* be the output of exp(−*ϵ*)-TGM returned by the database. The user uses (*π, ℓ*′) to obtain the loss-minimizing guess for the count, which is given by *T* (*z*; *π, ℓ*′, exp(−*ϵ*), *n*) (see Equation 1).

Finally, the user thresholds this number to obtain a binary answer for the membership query, given by **1**{*T*(*z*; *π, ℓ*′, exp(−*ϵ*), *n*) > 0}. Note that the application of *α*-TGM using our transformed loss *ℓ*′ is known to be optimal only given our generalized notion of utility we achieved in the previous section (Theorem 2); *ℓ*′(*c, y*) cannot be expressed as a monotonically increasing *f* (|*c* − *y*|) as required by Theorem 1, nor is it sufficient to drop only the symmetry assumption and consider quasiconvex *f* (*c* − *y*) with minimum at zero—we require the flexibility to set a different loss function *f*_*c*_(*c* − *y*) for each value of *c*.

## 4 Results

### 4.1 Overview of Our System

Here we describe the overall workflow of our optimal differential privacy mechanism for biomedical database queries (Figure 1). Suppose the database *D* consists of data from *n* individuals, *d*_1_, …, *d*_*n*_. First, the user chooses a desired privacy level, denoted by *ϵ*, and a query. For both cohort discovery and variant lookup scenarios, the query can be represented using a predicate *s*, which evaluates to true (1) or false (0) for each individual *d*_*i*_ in the database, indicating whether he or she matches the desired criteria for the cohort or has a variant of interest. The user is interested in 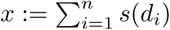 for cohort discovery and **1**{*x* > 0} for variant lookup, where **1**{·} is the indicator function. The user submits (*ϵ, s*) to the database server. Next, the server uses the truncated *α*-geometric mechanism (with the parameter *α* = exp(*ϵ*)) to release a differentially-private statistic 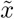 of *x* to the user. Then, the user chooses a prior distribution *π* over *x*, representing his or her belief about the underlying data distribution, and a loss function *ℓ*(*x, y*), representing how undesirable obtaining a query result *y* is, given *x*, where *y* ∈ {0, …, *n*} for cohort discovery and *y* ∈ {0, 1} for variant lookup. Based on the chosen *π* and *ℓ*, 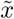 is mapped to a value *y*, which the user interprets as the answer to their query. Our scheme satisfies *ϵ*-differential privacy and provably minimizes the expected loss of the query result among all stochastic mechanisms that achieve *ϵ*-differential privacy, for both cohort discovery and variant lookup scenarios (Methods).

**Figure 1:**
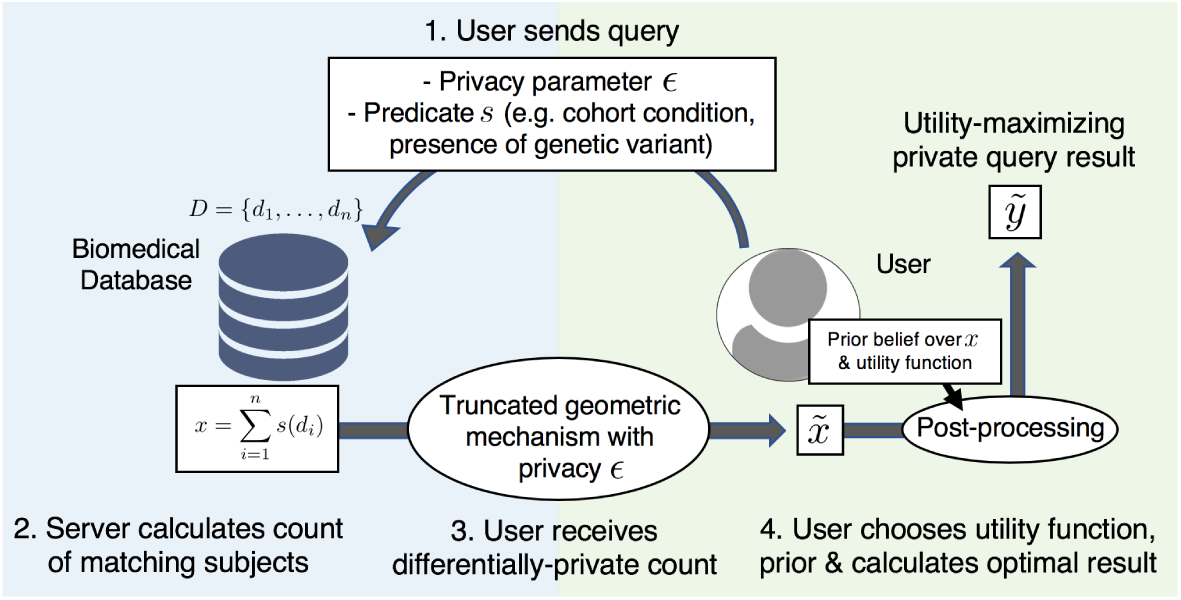
Overview of our optimal differential privacy schemes for biomedical queries. Given a count or membership query from the user, the database returns a differentially private count of individuals matching the query using the generic truncated *α*-geometric mechanism [16]. The user locally transforms the result based on his or her chosen the generic rior belief over the true count and a utility function. As we showed, the final result provably maximizes expected utility over all differential privacy mechanisms with the same privacy level.

Regarding the computational cost of our system, the overhead for the server is negligible as it randomly samples one additional number per query. Communication is identical to the underlying system without differential privacy, except for the inclusion of a privacy parameter *ϵ* in the query. Local post-processing by the user takes *O*(*n*^2^) time per query for a database of size *n* in general; however, the structure of loss functions in our application scenarios can be exploited to reduce the complexity to *O*(*n* log *n*) for cohort discovery and *O*(*n*) for variant lookup, based on fast Fourier transform (FFT)-based convolution and sparse matrix multiplication for computing Eq. 1, respectively. As a result, we achieve post-processing times of less than a second for both query types on a standard laptop computer even when *n* is a million.

### 4.2 Differentially-private cohort discovery with maximum utility

We first evaluated the utility of our proposed approach for cohort discovery. For baseline, we compared our approach to the exponential mechanism proposed by Vinterbo et al. [9], and the Laplace mechanism used in the MedCo framework of Raisaro et al. [13]. Note that the exponential mechanism takes a utility function as input, which we set to the negative of the loss function used in our mechanism. This linear mapping is motivated by the existing theory showing that exponential mechanism approximately maximizes the expected utility [26], which corresponds to minimizing the expected loss in our setting. In addition, because neither exponential nor Laplace mechanisms have a notion of prior distribution over the true count, we set our prior distribution to uniform, which represents a neutral prior.

As expected, our approach achieves the lowest expected loss across all values of privacy parameter *ϵ* for both symmetric and asymmetric loss functions (Figure 2). In fact, our theoretical results suggest any mechanism that achieves *ϵ*-differential privacy cannot perform better than our approach on average in both symmetric and asymmetric settings (Methods). Comparing the probability distribution over the query result, we see that our approach results in a more concentrated probability mass near the true count. It is worth noting that, with asymmetric loss, the mode of the distribution over the query result does not align with the true count, which is a result of the skewness of the loss function.

**Figure 2:**
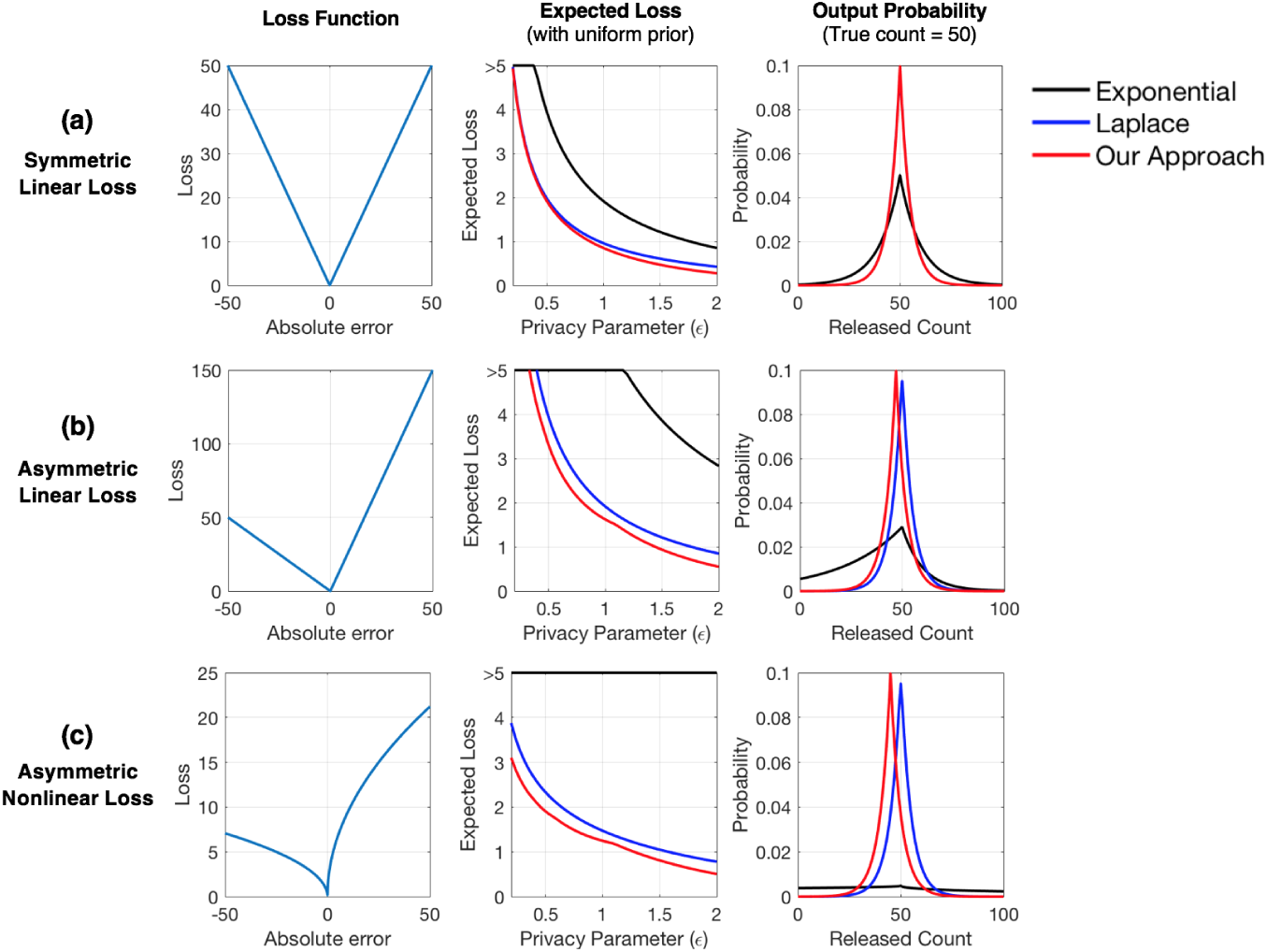
Our approach improves the utility of medical cohort discovery with differential privacy. We compared the performance of our optimal differential privacy mechanism for count queries to the exponential [9] and Laplace [13] mechanisms for different choices of loss functions (rows). We considered the following parameter settings for Vinterbo et al.’s loss function [9] described in Methods: (a) *α*_*−*_ = *α*_+_ = *β*_*−*_ = *β*_+_ = 1, (b) *α*_*−*_ = *α*_+_ = 1, *β*_*−*_ = 1, *β*_+_ = 2, and (c) *α*_*−*_ = *α*_+_ = 0.5, *β*_*−*_ = 1, *β*_+_ = 2. In each row, the subfigures show the shape of loss function (left), expected loss over a range of privacy parameters *E* (center), and a sample probability distribution over the private query result, with a true count of 50 and *E* = 0.2 (right). We used the uniform prior for our mechanism. Overall, our approach reduces the expected loss while maintaining the same level of privacy.

The fact that our margin of improvement over the Laplace mechanism is smaller than that over the exponential mechanism can be attributed to the fact that the Laplacian distribution, from which the perturbed count is sampled in the Laplace mechanism, closely approximates the geometric distribution used in our approach, which could be viewed as a discrete version of the Laplacian distribution. However, the Laplace mechanism is unable to tailor its answer to the user’s desired notion of utility (captured by the loss function) or prior belief over the data, resulting in a greater loss in utility in more sophisticated settings. For example, our experiments with asymmetric loss shows that the Laplace mechanism resorts to a suboptimal output distribution centered around the true count, leading to a more pronounced performance difference between our approach and the Laplace mechanism compared to the symmetric case (Figures 2b and c).

Importantly, a key aspect of our approach is that it allows the user to incorporate his or her belief about the underlying data distribution. For example, if a user is interested in the number of individuals with a disease that is known to be extremely rare, then it would be desirable to take advantage of this prior knowledge about the disease to further improve the accuracy of the query result, rather than assuming that every possible answer is equally likely (uniform prior). To demonstrate this capability, we performed an experiment where the query distribution is concentrated on small numbers (e.g. <5 out of 1000). We then evaluated the expected loss of our mechanism based on different choices of prior distributions of varying skewness, ranging from uniform to one that is highly skewed toward the low counts (Figure 3). Our results show that, indeed, adopting a prior distribution that is better aligned with the true result further reduces the expected loss of our system.

**Figure 3:**
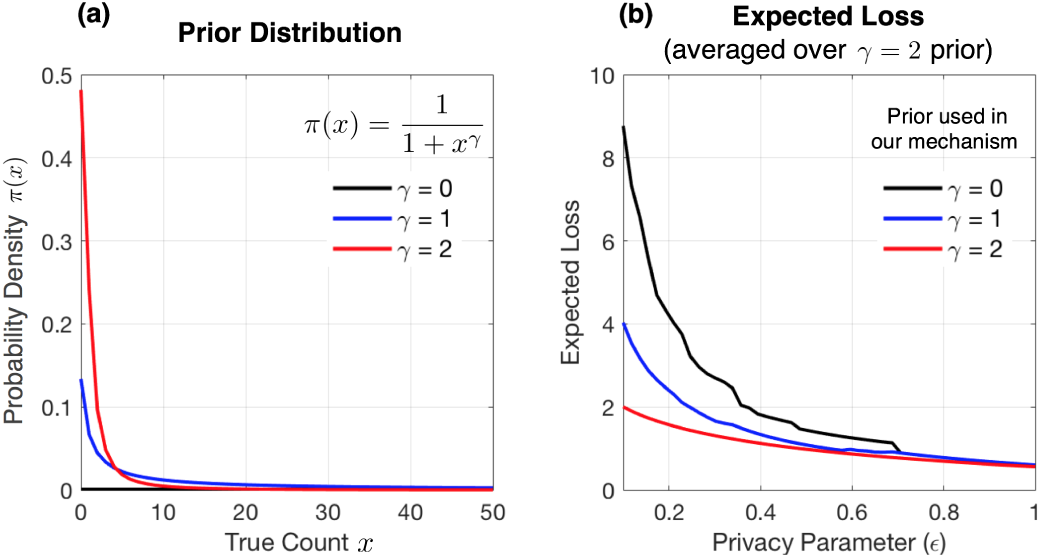
More accurate choice of prior distribution further improves the utility of private count query results. We show the effect of choosing different prior distributions over the true count (a) on the expected loss of count queries for different values of privacy parameters (b). We calculated the expected loss by averaging the results with respect to a highly-skewed prior distribution (*γ* = 2). As the prior distribution becomes more aligned with the true query distribution, the expected loss becomes smaller.

### 4.3 Differentially-private variant lookup with maximum utility

Variant lookup queries are becoming increasingly relevant for the biomedical community given the growing number of genomic beacon servers. Previously proposed privacy risk mitigation strategies for beacons [11] have aspects that are reminiscent of differential privacy (e.g. the notion of privacy budget), yet their privacy guarantees were based upon a simplified statistical model of genotype distributions in a population. As a result, the proposed strategies do not necessarily provide protection against attacks that take advantage of data patterns that lie outside of the model. For example, based on the 1000 Genomes Project dataset [27], we observed that selectively targeting genetic variants that deviate from Hardy-Weinberg equilibrium can lead to greater leakage of private information than captured by the previously-proposed privacy budget accounting scheme [11] (Supplementary Figure 1). Thus, theoretical frameworks such as differential privacy offer a valuable, more thorough alternative to mitigating privacy risks in beacon servers.

Our theoretical results enabled us to design an optimal differential privacy mechanism for variant lookup queries that maximizes a user-defined notion of utility for any desired privacy level (Methods). In our experiments, we consider two types of loss functions shown in Figure 4. *Linear* loss puts a linearly increasing penalty on answering that the variant is not found as its true count in the database grows (i.e., *ℓ*(*c*, 0) = *c* for *c* > 0). To additionally penalize false positive findings, we set the penalty for answering that the variant is found in the database when it is not (i.e., *ℓ*(0, 1)) to a higher value than one (in our case 100). *Uniform* loss equally penalizes all incorrect query answers regardless of how many times the query variant was observed in the database. Note that our framework allows users to choose their own loss function as long as it satisfies a straightforward assumption that the wrong answer is not preferred to the correct answer (Methods).

**Figure 4:**
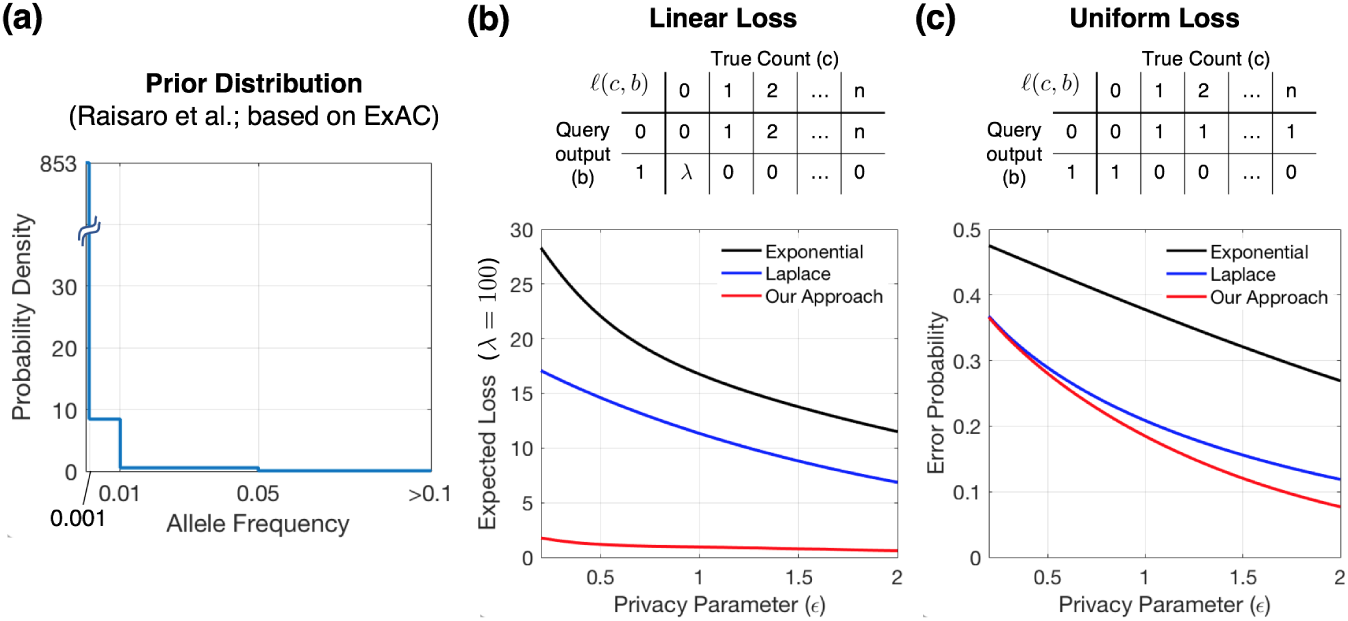
Our approach improves the utility of genomic variant lookup with differential privacy. Using the allele frequency distribution of variant lookup queries submitted to the ExAC database [11] as our prior distribution (a), we evaluated the expected loss of our optimal differential privacy mechanism for variant lookup with two different loss functions shown in tables: (b) *linear loss*, which employs a linearly increasing penalty for negatively answering the query as the number of observations in the database grows and an increased penalty for false positives parameterized by *λ*, and (c) *uniform loss*, where any error that results in a flipped query result incurs the same amount of penalty, in which case the expected loss can be interpretted as error probability. Overall, our approach achieves the lowest expected loss compared to existing differential privacy mechanisms.

To construct a realistic prior distribution on the true count *x*, we leverage the allele frequency (AF) breakdown of real-world variant lookup queries submitted to the ExAC browser [28] over a period of 12 weeks, provided by Raisaro et al. [11] (Figure 4a). Assuming a uniform distribution within each AF category defined by Raisaro et al., we transformed the prior over AF into a prior over the number of individuals in the database with the query variant, which we used both in our optimal mechanism and for averaging the results over the real-world query distribution.

We compared our approach to standard DP techniques, including exponential and Laplace mechanisms. For the exponential mechanism, we set the utility to be the negative of our loss function and release a binary answer directly based on the true count *x* of the query variant. For the Laplace mechanism, we first use it to obtain a privacy-preserving count 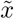 of *x*, then apply a threshold 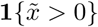 to obtain a binary answer. Both these approaches represent a straightforward application of the existing techniques.

Our results show a significant improvement in utility over existing mechanisms across a range of privacy parameters *ϵ*, with respect to linear loss (Figures 4b). In the case of uniform loss, our expected loss minimization problem reduces to minimization of the overall error probability of the system, since the loss function evaluates to one for the incorrect answer and zero for the correct answer in all cases. Under this setting, our mechanism also achieves smaller error probabilities compared to the existing differential privacy techniques (Figure 4c).

### 4.4 Differentially-private association tests with improved accuracy

Count queries used in cohort discovery represent a fundamental operation that can enable a range of down-stream analyses in a privacy-preserving manner, including association tests and histogram visualizations. Given that our optimal differential privacy framework improves the accuracy of count queries while achieving the same level of privacy, it is expected that these improvements will also translate into accuracy improvements in downstream tasks. Here, we set out to empirically demonstrate this idea using the *𝒳*^2^ association test as an example, which is a foundational technique for testing independence between pairs of categorical variables (e.g. clinical, genomic, and demographic characteristics of individuals).

To this end, we generated a large collection of sample 2-by-2 contingency tables with varying degrees of dependence between the two binary variables (one represented by the rows, and the other by the columns). Assuming these tables as ground truth, we then obtained a differentially private version of each table by separately invoking differential private count query on each of the four cells with privacy parameter *ϵ*. (Note that due to the parallel composition property of differential privacy (Methods), this overall procedure satisfies *ϵ*-DP, instead of 4*ϵ*.) We then computed the *𝒳*^2^ association statistic based on both the true table and the perturbed table for comparison. As expected, we observed that the differentially private *𝒳*^2^ statistics calculated based on our optimal mechanism more accurately matched the ground truth as opposed to other baseline mechanisms (Figure 5), though the Laplace mechanism achieves comparable performance due to its similarity to our mechanism under symmetric linear loss we adopted in this experiment. These results illustrate the broad impact of our approach beyond cohort discovery and variant lookup workflows. It would be interesting to see how these count query-based approaches compare to other DP mechanisms tailored for chi-squared tests [29] and whether one can design an optimal mechanism for calculating chi-squared statistics based on our techniques.

**Figure 5:**
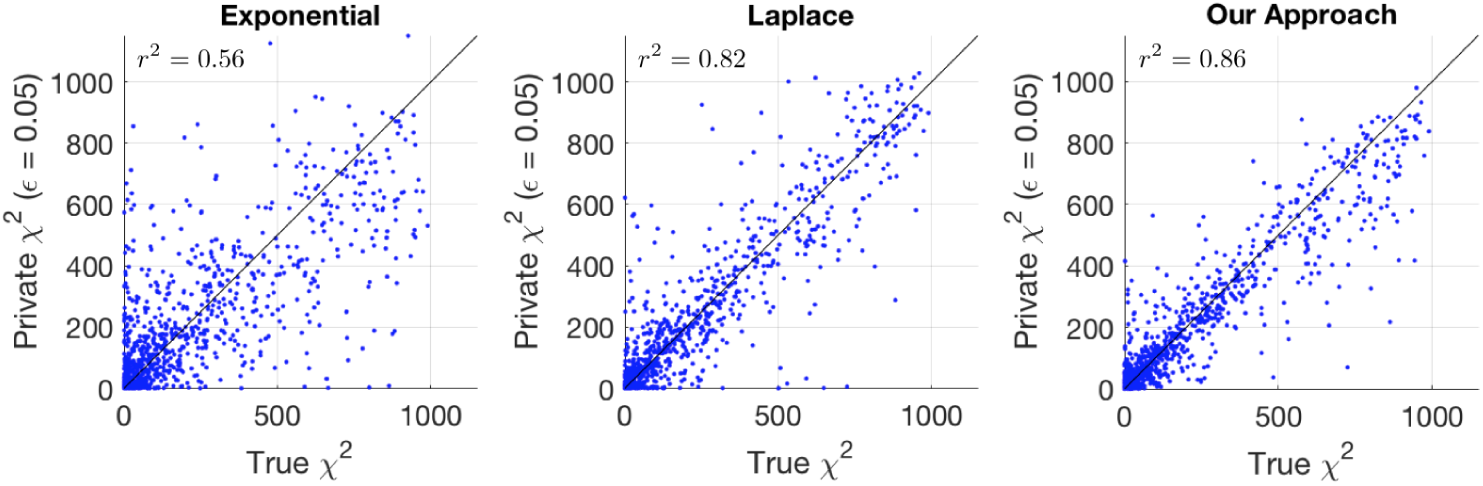
Our approach improves the accuracy of chi-squared association tests with differential privacy. We simulated datasets each including 1000 paired observations of two binary random variables with varying strengths of association. We considered a differentially private (DP) scheme for releasing the chi-squared association statistic between the two variables for each dataset, where we use DP count queries to construct a 2 × 2 contingency table, based on which the statistic is computed as post-processing. The subfigures show the agreement between the DP statistics and the ground truth for different choices of DP count query mechanisms. We optimized our mechanism with respect to uniform prior and a symmetric linear loss function. Our approach achieves the best accuracy overall, comparable to the Laplace mechanism yet considerably more accurate than the exponential mechanism.

## 5 Discussion

We presented optimal differential privacy (DP) mechanisms for a range of biomedical database queries, including medical cohort discovery and genomic variant lookup workflows. We achieved these results by newly extending the well-studied *α*-geometric mechanism of Ghosh et al. [16] to a broader class of utility functions for count queries and to the problem of membership queries. Furthermore, we empirically demonstrated that our approach improves the accuracy of query responses over existing DP mechanisms while achieving the same level of privacy. Given the optimality of our schemes, *our results illustrate the theoretical boundaries of leveraging DP for privacy risk mitigation in biomedical query-answering systems.*

It should be noted that due to the stringent nature of DP, the smallest possible error probabilities for beacons achieved by our optimal mechanism could still be too high for real-world systems to endure without significantly restricting their use. This limitation primarily arises from the fact that the queries submitted to beacons are highly skewed toward *rare* variants, which are also the most sensitive data from the perspective of DP, thus inflating the overall error probability. Our optimality proofs suggest that in order to surmount this challenge, we will require fundamental shifts in theory to expand the scope of DP to allow more permissive frameworks that remain useful in practical scenarios. Another solution may be hybrid systems that combine DP with traditional access control procedures to allow the sharing of highly-sensitive query results based on trust, while securing and facilitating other types of queries that are more amenable to DP. In particular, our DP techniques achieve high accuracy for count query results that are sufficiently large (e.g. greater than 50) and membership queries for which the underlying count is similarly large; both cases will become increasingly common as biomedical databases grow in size.

There are several interesting methodological directions that merit further research. It may be possible to design optimal DP mechanisms for more complex tasks beyond count and membership queries, including emerging applications of DP in federated machine learning [17]. We also plan to explore better ways to address the setting where each user submits a potentially large number of queries; although current work considers each query independently, one could exploit the structure of common biomedical queries to develop more effective methods to compose multiple queries, such that the overall privacy cost is smaller than that obtained by general-purpose composition techniques [21]. Lastly, in line with the work of Raisaro et al. [13], we need further efforts to incorporate DP mechanisms into federated systems that enable collaborative analysis across a group of entities with isolated datasets, who are unable to directly pool the data due to regulatory constraints or conflicting interests (e.g. [30,31]). Together, these efforts will help us bring cutting-edge advances in differential privacy to real-world biomedical data sharing platforms in order to empower researchers while enhancing protection for individuals.

## Appendix

**Figure S1:**
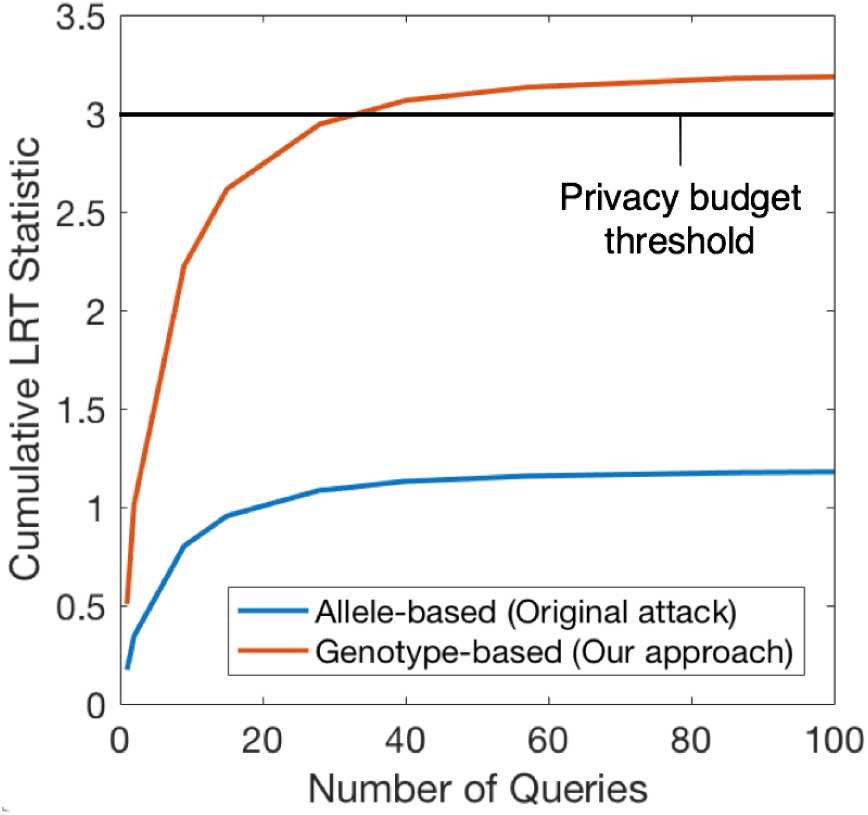
Taking advantage of model assumption violations to bypass existing privacy risk mitigation strategies for genomic beacons. Raisaro et al. [11] proposed to quantify the privacy leakage of variant lookup queries by calculating the likelihood ratio between the null hypothesis (target individual is not in the database) and the alternative hypothesis (target individual is in the database). In either setting, the data likelihood is calculated based on a simple model of genotype distribution, where each of the two alleles in an individual is sampled from a Bernoulli distribution whose parameter is set to the allele frequency in a corresponding population. More precisely, Raisaro et al. calculate the risk of each query as − log(1 − *D*_*n*_) where *D*_*n*_ represents the probability of the query variant not being present in a database of size *n* under the null hypothesis. *D*_*n*_ is computed as (1 − *f*)^2*n*^ given the Bernoulli model and an estimated frequency of the alternative allele *f*. However, for variants that deviate from the Hardy-Weinberg equilibrium (HWE), this calculation does not result in an accurate estimate of the risk. Here we considered a more refined model of the genotype distribution, which directly estimates the frequencies of homozygous reference, homozygous alternative, and heterozygous genotypes, then calculates *D*_*n*_ instead as *D*_*n*_ = *h*^*n*^, where *h* denotes the frequency of the homozygous reference genotype. Note that this reduces to Raisaro et al.’s approach under HWE, since *h* = (1 − *f*)^2^. We tested this approach on the 1000 Genomes Project dataset, which includes 2504 individuals. We randomly chose half of the individuals to estimate the genotype frequencies. Among the remaining half, we held out 100 individuals for testing purposes and constructed a beacon from the remaining individuals. The figure shows the cumulative risk (i.e. the summation of − log(1 − *D*_*n*_) over the query history) with one of the test subjects as a target, under the scenario where the attacker queries variants with the largest departure from the HWE first. Our more accurate estimation of the risk based on genotype frequencies (in red) reveals considerably greater privacy leakage than captured by the previous approach (in blue). In fact, in our analysis the cumulative risk exceeds the privacy budget threshold given by Raisaro et al. (− log(0.05); black horizontal line) in less than 40 queries, which the original scheme fails to detect. LRT: Likelihood ratio test.

### Proof of Theorem 2: Optimality of truncated *α*-geometric mechanism for count query with more general loss functions

Here, we answer the open question posed in Ghosh et al. about the optimality of *α*-geometric mechanism for asymmetric loss functions in the affirmative. In addition, we further generalize this result to loss functions that are separately defined for each possible value of the true count. The only step in the original proof that relies on the properties of the loss function is in the following lemma. Thus, it is sufficient to prove that this lemma holds for more general loss functions.

#### Lemma

(Ghosh et al., Lemma 5.5). For every user *u* with a full-support prior and a strictly legal loss function, every optimal direct mechanism for *u* has a unimodal signature.

Note that Ghosh et al. define (strictly) *legal* loss function *ℓ*(*i, j*) for input *i* ∈ {0, …, *n*} and output *j* ∈ {0, …, *n*} as a (strictly) monotone increasing function that depends only on |*i* − *j*|. Below we prove that the above lemma still holds even if we expand the scope of legal loss functions to those that can be represented using a pair of monotone increasing functions 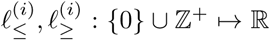 for every *i* ∈ {0, …, *n*}, whose values coincide at zero (i.e., 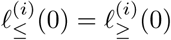, and

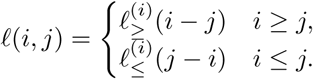

In other words, this expanded class of loss functions include any function that monotonically increases in value as we change the output *j* in either direction starting from the true count *i, without* requiring that it is (i) symmetric 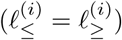 or (ii) identically shaped for different values of *i* (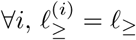 and 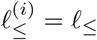 for some *ℓ*_≥_, *ℓ*_≤_). Note that for the proof of the above lemma it is acceptable to assume *strict* monotonicity of loss functions; Ghosh et al. presents a limiting argument to discharge this assumption later in their proof, which applies analogously to our generalized loss function.

#### Lemma.

For every user *u* with a full-support prior and a loss function who is legal by our generalized definition, every optimal direct mechanism for *u* has a unimodal signature.

The proof of the above lemma using our generalized definition of legal loss function is as follows. Given a full-support prior *π* ≻ 0 and strictly legal loss function 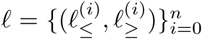 for a database of size *n*, let *X* be an optimal direct mechanism with signature Σ (where Σ is the constraint matrix from Ghosh et al.). As in the original proof, we want to show that Σ is unimodal: that is to say each row begins with some number of ↓ entries (possibly 0), followed by zero or one S entries, followed by some number of ↑ entries (possibly 0). In order to prove this, it suffices to show that there is no row *h* and columns *k* < *m* such that *σ*_*hk*_ ∈ {*S*, ↑} and *σ*_*hm*_ ∈ {*S*, ↓}.

Assume there exist such *h, k*, and *m*. We first show that *k* < *h* < *m* by deriving contradiction from *h* ≤ *k* or *h* ≥ *m*. Assume *h* ≤ *k* (the *m* ≤ *h* case follows an analogous argument). The facts that *σ*_*hk*_ ∈ {*S*, ↑} and *σ*_*hm*_ ∈ {*S*, ↓} imply the following inequalities:

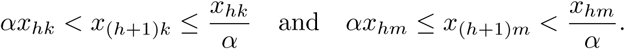

Following the same argument given in the original proof, this means that we can find a small *λ* > 0 such that defining a revised mechanism *X*′ where 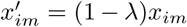 and 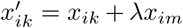 for all *i* ∈ {0, 1, …, *h*}, while keeping the remaining entries the same as *X*, results in a feasible mechanism for the given problem. Note that the difference in expected loss between the two mechanisms is given as

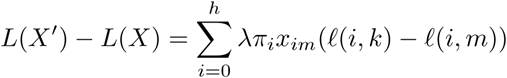

Given that *h* ≤ *k* and the initial assumption *k* < *m*, we have

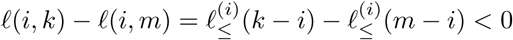

for all 0 ≤ *i* ≤ *h*, using the strict monotonicity of 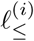. Thus, *X*′ achieves a strictly lower expected loss than *X* (given *λ* > 0 and *π* ≻ 0), which is a contradiction. An analogous line of reasoning, which instead involves moving the probability mass in rows *i* ∈ {*h* +1, …, *n*} of *X* and the monotonicity of 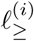, shows contradiction for *h* ≥ *m*. Thus, *k* < *h* < *m*.

Another implication of the above result is that for any tuple (*h*′, *k*′, *m*′) where *k*′ < *m*′, and *h*′ ≤ *k*′ or *h*′ ≥ *m*′, either *σ*_*h*′*k*′_ =↓ or *σ*_*h*′ *m*′_ =↑. We now use this fact to derive contradiction for the case where *k* < *h* < *m*. Our assumption states that *σ*_*hk*_ ∈ {*S*, ↑} and *σ*_*hm*_ ∈ {*S*, ↓}. Applying the above logic to tuples (*h, k, h*) and (*h, h, m*), we obtain that *σ*_*hh*_ = ↑ from the former tuple, and *σ*_*hh*_ = ↓ from the latter tuple, which together gives a contradiction. Thus, all possible values of *h* result in a contradiction, proving the original statement of the lemma.

### Proof of Theorem 3: Optimality of truncated *α*-geometric mechanism for membership query

Let *c* ∈ {0, …, *n*} be the number of individuals in a database matching the predicate of a given membership query (e.g. presence of a genetic variant). Let *π*(*c*) be a user-defined prior belief over *c*, and *ℓ*(*c, b*) be a user-defined loss function representing the disutility of receiving a membership query result *b* ∈ {0, 1} when the true count is *c*. As stated in the theorem, assume *ℓ* satisfies *ℓ*(0, 0) ≤ *ℓ*(0, 1) and *ℓ*(*c*, 0) ≥ *ℓ*(*c*, 1) for *c* > 0.

The expected loss minimization problem for *ϵ*-differentially private membership query, parameterized by a conditional probability distribution *q*(*b*|*c*), can be expressed as follows.

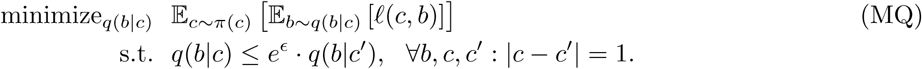

We want to show that an optimal solution *q*(*b*|*c*) for the above problem is given by the exp(−*ϵ*)-TGM with a (*π, ℓ*)-dependent post-processing.

Now consider a transformed loss function *ℓ*′, defined as

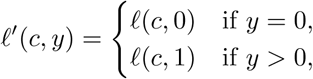

for *c, y* ∈ {0, …, *n*}. First, note that this loss function satisfies the conditions of Theorem 2; for each value of *c*, we have that *ℓ*′^(*i*)^(*y* − *i*) := *ℓ*′(*i, y*) is monotone increasing in both directions from zero, given our initial assumptions about *ℓ*. Therefore, we can invoke Theorem 2 using (*π, ℓ*′) to obtain that there is a (*π, ℓ*′)-dependent post-processing *T* such that, given the output *z* of a differentially private release of the true count *c* based on the exp(−*ϵ*)-TGM, *T* (*z*) represents the output of an optimal *ϵ*-DP scheme that is an optimal solution to the following count query problem.

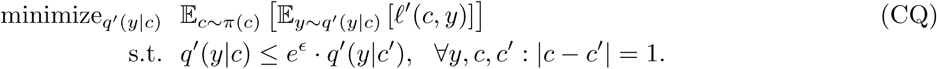

We next prove that the problems MQ and CQ are in fact interchangeable. First, note that any feasible solution *q*′ of CQ, can be mapped to a feasible solution *q* of MQ by setting

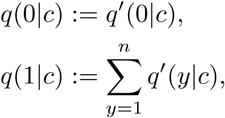

which is equivalent to post-processing *y* as *b* = **1** {*y* > 0}. If *y* is *ϵ*-DP, then we have that *b* is also *ϵ*-DP, which implies feasibility of the mapped *q*. Furthermore, because *ℓ*′(*c, y*) = *ℓ*(*c*, 1) for all *y* > 0, we have that (*)

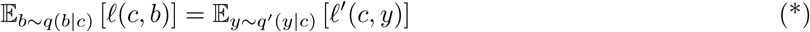

for our construction of *q* based on *q*′. Therefore, for any feasible solution of CQ, a corresponding solution exists for MQ that achieves the same objective value.

Next, we consider the reverse direction. Given any feasible solution *q* of MQ, let *q*′ be a solution for CQ constructed as follows:

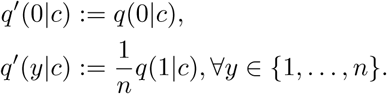

This is akin to post-processing *b* by setting *y* = 0 if *b* = 0, and *y* ∼ Unif({1, …, *n*}) if *b* = 1. Therefore, if *b* is *ϵ*-DP then *y* is also *ϵ*-DP, proving that the above *q*′ is a feasible solution for CQ. Because 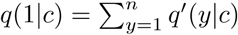 by construction, (*) still holds. This proves that for any feasible solution for MQ, there exists a feasible solution for CQ with the same objective value.

Now putting the two directions together, we have that an optimal solution for CQ, represented by *T* (*z*), can be post-processed as **1** {*T* (*z*) > 0} to obtain an optimal solution for MQ. This is because, if we assume a better solution for MQ exists, then we can map it to CQ to obtain a better solution than *T* (*z*), which is a contradiction. Note that **1** {*T* (*z*) > 0} can be viewed as a post-processing of the exp(−*ϵ*)-TGM output *z*, where *T* depends on the user-provided *π* and *ℓ*. This concludes the proof of the theorem. □

### Optimal choice of *ω* in exponential mechanism

#### Algorithm 1 Out modified exponential mechanism for count queries

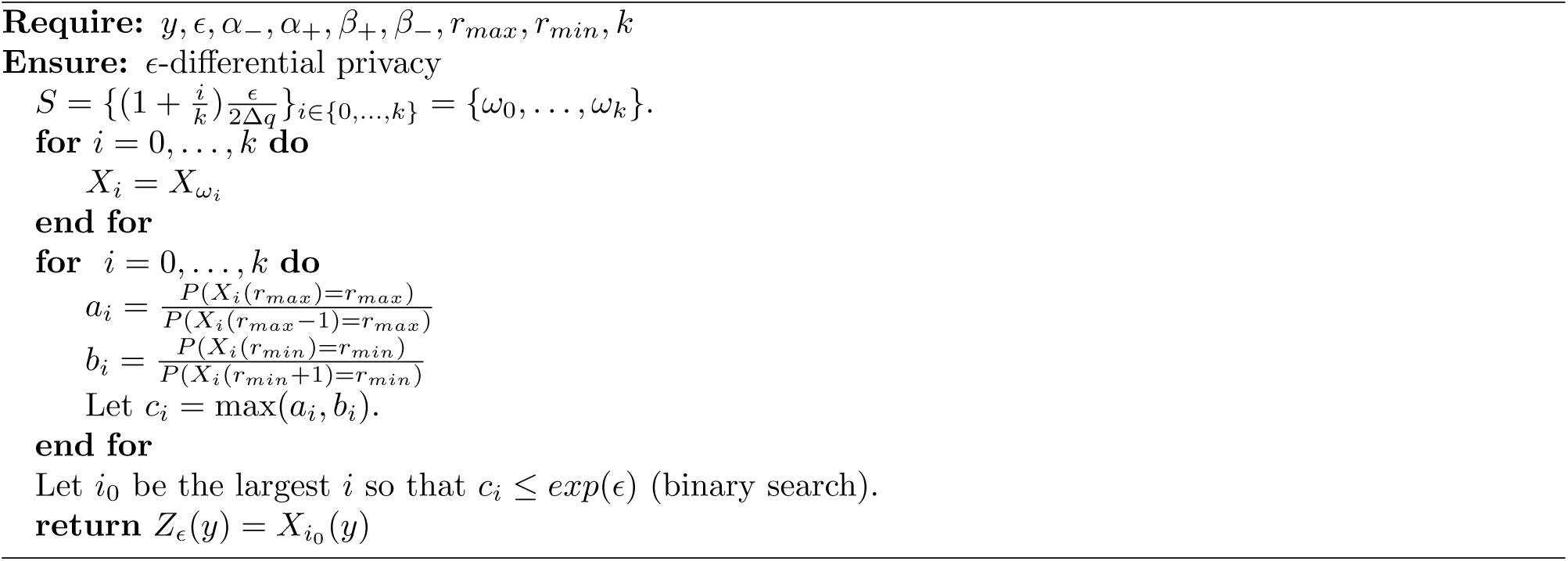

To prove this algorithm is differentially private, we first need to prove the following:

#### Theorem 4.

*Assume that α*_+_, *α*_−_ ∈ [0, 1], *β*_+_ > 0, *β*_−_ > 0, *and that*

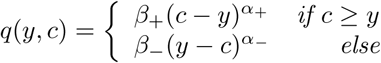

*Let X*(*y*) *be a random variable defined so that P*(*X*(*y*) = *c*) *is proportional to exp*(−*ωq*(*y, c*)) *for all integers c* ∈ [*r*_*min*_, *r*_*max*_] *and y* ∈ [0, *n*]. *Then we have that, if*

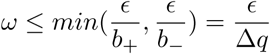

*P*(*X*(*r*_*max*_) = *r*_*max*_) ≤ *exp*(*ϵ*)*P*(*X*(*r*_*max*_−1) = *r*_*max*_) *and P*(*X*(*r*_*min*_) = *r*_*min*_) ≤ *exp*(*ϵ*)*P*(*X*(*r*_*min*_+1) = *r*_*min*_) *then X*(*y*) *is ϵ-differentially private.*

*Proof.* Let 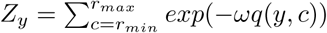 then by definition

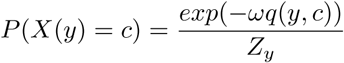

For a given *c* we are interested in 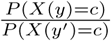 and 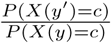 where *y* = *y*′ − 1.

There are three cases: the first when *y* ≥ *r*_*max*_, the second when *y* < *r*_*min*_, and the third when *y* ∈ [*r*_*min*_, *r*_*max*_ − 1].

Consider the case when *y* < *r*_*min*_. Then *Z*_*y*_ < *Z*_*y*+1_ so

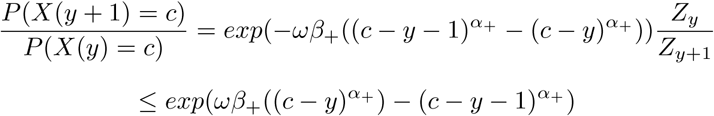

Since *α*_+_ ≤ 1, *c* − *y* > *c* − *y* − 1 ≥ 0 we see that 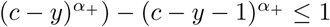 so the above is

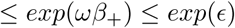

just as we wanted. On the other hand consider

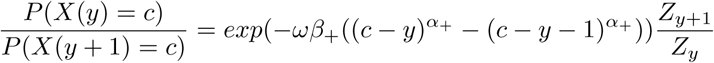

Note that 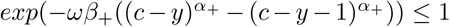 so the above is 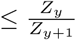. Since *q* has sensitivity bounded by *max*(*b*_−_, *b*_+_) we see that 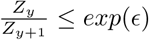, so putting the above together

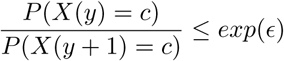

By a similar argument, when *y* ≥ *r*_*max*_ we have that

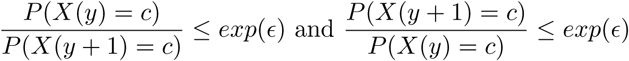

Finally consider the case when *r*_*min*_ ≤ *y* < *r*_*max*_. Then note that

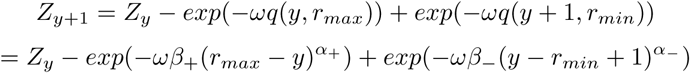

We will consider 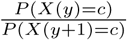. If *c* ≥ *y* then *q*(*y, c*) > *q*(*y* + 1, *c*) so since 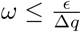 we get

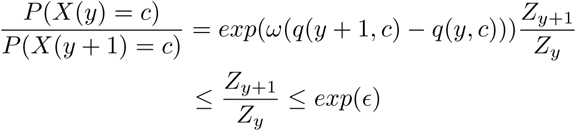

Therefore assume that *c* > *y*. Then *exp*(*ω*(*q*(*y* +1, *c*)−*q*(*y, c*))) ≤ *exp*(*ωβ*_+_), which is achieved when *c* = *y* +1. We next consider 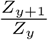 Note that 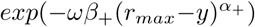 is increasing in *y* while 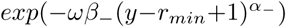 is decreasing in *y*. Thus it is easy to see that there exists a *y*_0_ ∈ (*r*_*min*_, *r*_*max*_) where 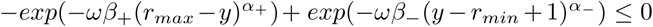 if *y* ≥ *y*_0_ while 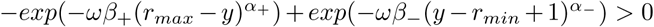 if *y* < *y*_0_.

Thus if *y* ≥ *y*_0_ we see that 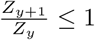 so

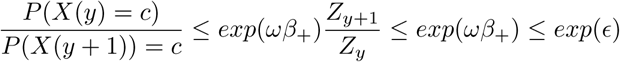

If, on the other hand, *y* < *y*_0_ this implies that *Z*_*y*_ < *Z*_*y*+1_, so 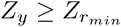, while

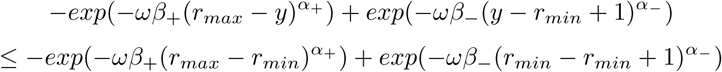

so

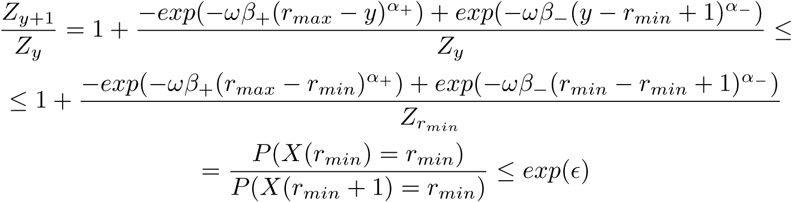

Thus *P* (*X*(*y*) = *c*) ≤ *exp*(*ϵ*)*P* (*X*(*y* + 1) = *c*) for all *y* and *c*. A symmetric argument shows that

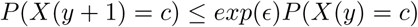

So *X* is *ϵ*-differentially private, just as we wanted. □

#### Corollary 4.1.

*The modified exponential method for count queries (Algorithm 1) is ϵ-differentially private.*

If we fix *k* and let *ω*_*ϵ*_ be the *ω* parameter chosen by Algorithm 1 we get that (since 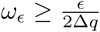):

#### Corollary 4.2.

*For a given ϵ and c we get that*

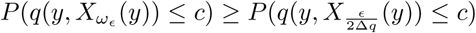

This algorithm was implemented in python and the code is available at https://github.com/seanken/DP_count.

